# A neural theory for counting memories

**DOI:** 10.1101/2022.05.18.492502

**Authors:** Sanjoy Dasgupta, Daisuke Hattori, Saket Navlakha

## Abstract

“I’ve never smelled anything like this.” “I’ve seen you once before.” “I’ve heard this song many times.” Keeping track of the number of times different stimuli have been experienced is a critical computation for behavior. This computation occurs ubiquitously across sensory modalities, and naturally without reward or punishment. However, the neural circuitry that mediates this computation remains unknown. Here, we propose a theoretical two-layer neural circuit that can store counts of stimulus occurrence frequencies. This circuit implements a data structure, called a *count sketch*, that is commonly used in computer science to maintain item frequencies in streaming data. Our first model implements the count sketch data structure using Hebbian synapses and outputs stimulus-specific frequencies. Our second model uses anti-Hebbian plasticity and only tracks frequencies within four count categories (“1-2-3-many”), which we suggest makes a better trade-off between the number of categories that need to be distinguished and the potential ethological value of those categories. Using real-world datasets, we show how both models can closely track the frequencies of different stimuli experienced, while being robust to noise, thus expanding the traditional novelty-familiarity memory axis from binary to continuous. Finally, we show that an implementation of the “1-2-3-many” count sketch — including network architecture, synaptic plasticity rule, and output neuron that encodes count categories — exists in a novelty detection circuit in the insect mushroom body, and we argue that similar circuit motifs also appear in mammals, suggesting that basic memory counting machinery may be broadly conserved.

## Introduction

Estimating the frequencies of different stimuli experienced (e.g., odors, faces, sounds) is an important computation that requires storing and updating the number of times each stimulus has been observed. This computation occurs naturally, without effort, and allows organisms to make rapid behavioral decisions, absent any specific details about the memory [1].

One line of evidence that the brain keeps track of stimulus occurrence frequencies comes from studies of recognition memory [2], which report neurons whose activity encodes whether a stimulus is novel or familiar. Recognition memory exists for many types of stimuli, including visual [3], auditory [4–6], and olfactory [7, 8]. For example, when monkeys are shown a sequence of pictures, neurons in the perirhinal cortex exhibit responses that relate to whether the image was previously shown [9]. Most studies report neurons whose response magnitudes decrease with familiarity; i.e., neurons show strong responses upon the first presentation of the stimulus, and weaker responses to subsequent presentations (called repetition suppression [10]). Others have found neurons that become more active with familiarity (called repetition enhancement [11, 12]). While many computational models of recognition memory have been proposed [13–17] (see review by Bogacz and Brown [18]), most models consider familiarity discrimination as a binary problem — is the stimulus novel or familiar? — as opposed to a continuous problem, where the desired output is an estimate of how many times the stimulus has been experienced. In addition, classic models are not well integrated with modern experimental data revealing how neural circuits represent stimuli in high-dimensional spaces and update their frequencies at synaptic resolution.

Frequency estimation is distinct from the numbers sense [19, 20], which underlies the ability to perform approximate numerical comparisons. For example, when frogs chose between patches of food items, their choice between three and four items is random, but they reliably chose six items over three [20]. Similar behaviors have been observed across the animal kingdom [21] — including in primates [22, 23], reptiles [24], fish [25–27], birds [28], flies [29], and bees [30–32] — without relying on language or numerical symbols. In monkeys and humans, there is evidence of neurons that encode numbers [33, 34]; number neurons respond most strongly to their preferred numbers, but they also respond to a lesser extent to adjacent numbers, and thus they have a bell-shaped response function [35, 36]. While useful for quantifying magnitudes — the number of food items in a patch, the number of predators in a group — the numbers sense does not provide a way to store a mapping from observed items to frequency counts, nor a way to update counts as items are experienced over time.

In computer science, frequency estimation comes up in many applications, such as keeping track of the number of times different videos are watched or different songs are played, to identify popular content. This problem is commonly solved using a data structure called a *count sketch* [37, 38]. Much like how an artistic sketch provides a quick approximation of a complex drawing, a “sketch” is a data structure that provides approximate answers to a query, while consuming substantially (often exponentially) less space than what would be required to store all of the data. A “count sketch” is a sketch that supports the frequency estimation query; i.e., “how many times have I seen item *x*?”. Count sketches are primarily used in instances where large amounts of data are continuously processed and where storing all of the data is prohibitive.

Here, we develop a theory for keeping track of stimulus occurrence frequencies, while being tolerant to noise. Our proposed neural circuit implements a count sketch using a staple two-layer neural architecture: a sparse, high-dimensional stimulus encoding layer that synapses onto a decoding layer with one neuron, which outputs the frequency of any observed stimulus. We also propose a variant of the model, called the “1-2-3-many” sketch, that only tracks frequencies within four categories, ranging from novel (frequency = 1) to very familiar (frequency > 3). Both models effectively expand the classic novelty-familiarity axis from a binary state memory system to a continuous one. We empirically demonstrate the accuracy of both neural count sketches on three datasets, and we derive mathematical bounds of their error as a function of environmental and neural variables (e.g., number of stimuli observed, number of encoding neurons, synaptic precision). Finally, we show that all the hardware needed to implement the “1-2-3-many” count sketch exists in the insect mushroom body, and re-analysis of published experimental data indeed shows that novelty responses can be distinguished along the four categories proposed. We conclude by raising several testable experimental hypotheses, and by describing other brain regions that have all the machinery needed to support memory counting.

## Results

We begin by presenting the count sketch data structure as a solution to the memory counting problem. We then present a neural implementation of the count sketch and show that it works well in practice and in theory. Finally, we show that three main requirements of our model — the circuit architecture, the synaptic plasticity rule induced after stimulus observation, and the response precision of the counting neuron — exist in the insect mushroom body.

### The count sketch data structure for frequency estimation in streaming data

Say we are given a sequence of observed items, where each item is drawn from a set 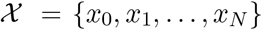 of *N* possible items. The sequence can contain the same item multiple times, and we would like to keep track of the number of times each unique item is seen. A hash table mapping keys (items) to values (counts) would provide exact counts but would require storing each item in its entirety, which would be very costly if the items are large (e.g., videos or songs) and numerous. A *count sketch* is a data structure that provides approximate counts, while only requiring a few bits of storage space per item, no matter how big the items themselves are.

A count sketch stores a frequency table for items using a 2D matrix *C* with *k* rows and *v* columns (Figure 1A). Each row is associated with a hash function *h* : *x* → [*v*]; i.e., the function takes as input some item *x* and maps it to a column index in *C*. The *k* hash functions are pairwise independent and random. For example, in Figure 1A, there are three hash functions (*k* = 3). Each entry in *C* corresponds to a counter and is initialized to 0.

**Figure 1:**
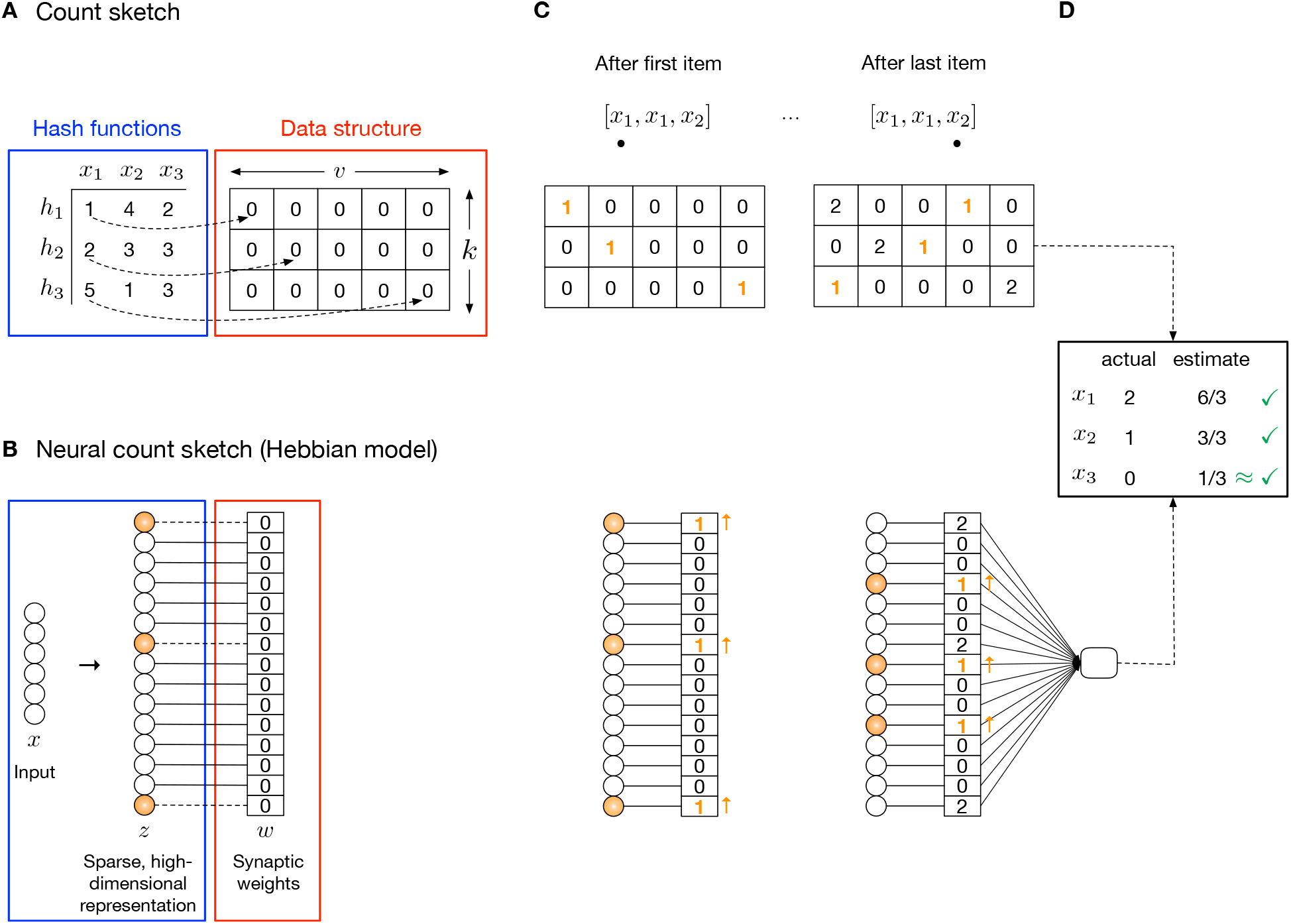
The count sketch and corresponding neural circuit implementation. **A)** The count sketch data structure is a 2D matrix *C* of counters with *k* rows and *v* columns. There is one hash function *h* per row, each of which determines which column in the row is modified when an item *x* is observed (dotted arrows). **B)** The neural implementation of a count sketch uses a 1D array of *k* × *v* synapses. When an item is observed, the synapses of the *k* pre-synaptic neurons that are active for the item (orange highlight) are modified. In this example, each item *x* is a *d*-dimensional vector. **C)** To insert an item from the sequence into the count sketch (top), *k* counters are incremented. For example, after the first time (*x*_1_) is observed, in the first row, the counter in the 1st column (i.e., *C*[1, 1]) is incremented by 1 since *h*_1_(*x*_1_) = 1. In the second row, *C*[2, 2] is incremented by 1 since *h*_2_(*x*_1_) = 2. In the third row, *C*[3, 5] is incremented by 1 since *h*_3_(*x*_1_) = 5. Similarly, in the Hebbian neural count sketch (bottom), the synaptic weights of the *k* activated pre-synaptic neurons for *x*_1_ (orange highlight) are incremented. In the anti-Hebbian model, all synaptic weights are initialized to 1, and synapses active for the item are decremented with each observation. **D)** To output an estimate of the frequency of item *x*_1_, the count sketch computes: 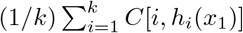 — i.e., the average of the predicted counts over the rows. This results in correct estimates for *x*_1_ and *x*_2_, and a near-correct estimate for *x*_3_. Similarly, the neural count sketch outputs: 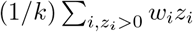 — i.e., the average of the weights of the activated neurons for the item.

To insert an item *x* into the count sketch, for each hash function *i*, we compute *j* = *h_i_*(*x*), and then we increment *C*[*i, j*] by 1. In Figure 1A, *h*_1_(*x*_1_) = 1, which means that the first hash function maps input *x*_1_ to column 1. So, when *x*_1_ is observed (Figure 1C, left), we increment *C*[1, 1] by 1. Similarly, *h*_2_(*x*_1_) = 2, which means we increment *C*[2, 2] by 1, and *h*_3_(*x*_1_) = 5, which means we increment *C*[3, 5] by 1. After these three entries are modified, we are finished inserting *x*_1_. This process repeats for each subsequent item (Figure 1C, right).

At any point, we can query the count sketch for the estimated frequency of item *x* (Figure 1D):

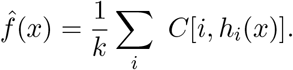

Intuitively, each row stores a predicted count for the item using a single hash function, which is then aggregated (averaged) over the rows into a final estimate. Other aggregate functions include median [37] and min [38–40], amongst others [41].

The accuracy of the estimate depends on the values chosen for *k* (the number of rows) and *v* (the number of columns). If *v* is large enough such that each unique item observed is mapped to a unique column index, then only a single row (*k* = 1) is needed to generate exact count estimates. However, in practice, hash collisions (overlaps) are likely, where a hash function maps two different items to the same column index. For example, in Figure 1D, the counts for *x*_1_ and *x*_2_ are exactly correct because each item is mapped to a unique set of column indices that do not overlap with those of other observed items. On the other hand, despite *x*_3_ never being observed in the input sequence, the count sketch would estimate its frequency to be 1/3 because *h*_2_ maps both *x*_3_ and *x*_2_ to the same column index (3). Thus, the level of approximation relates to the amount of overlap with other items, as well as the number of rows that are averaged over. Overall, larger values of *k* and *v* provide more accurate estimates, at the expense of larger space consumption.

### A neural implementation of a count sketch

There is a very simple way that neural circuits can implement a count sketch data structure (Figure 1B). The main idea is to “flatten” the 2D matrix of counters with *k* rows and *v* columns into a 1D array of *k* × *v* synapses. In the count sketch, each input modifies the values of *k* entries in the table (one per row). In the neural version, each input will modify *k* synaptic weights. The identity of these *k* synapses will be determined by a neural hash function, which will encode inputs using sparse, high-dimensional representations. Specifically, of the *k* × *v* pre-synaptic neurons, only *k* ≪ *v* will fire per input, and the synapses of these neurons are modified for the input. Post-synaptically, there is one decoding neuron that reads-out from the encoding neurons and outputs a frequency for the given stimulus.

These three pieces (stimulus encoding, synapse weight updating, and frequency decoding) are described below.

#### Stimulus encoding

The first piece determines which pre-synaptic neurons are active for an input. This requires designing a neural hash function, 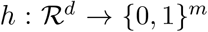, which takes some input vector 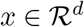 and assigns it to a point in m-dimensional space, where *m* = *kv*. A canonical way to do this is via random projection and sparsification [42]. This motif is used widely, including in the olfactory system [43–45], hippocampus [46], and cerebellum [47], to create sparse, high-dimensional representations for inputs [48, 49].

In the random projection step, we compute 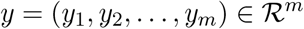 by:

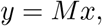

where *M* is a random matrix of size *m* × *d*. For example, *M* can be a Gaussian random matrix, where each value is drawn i.i.d. from 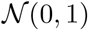; or, it could be a sparse binary matrix, where each row of *M* has a small number of 1s and the rest of the values are 0.

In the sparsification step, we compute *z* = (*z*_1_, *z*_2_, …, *z_m_*) ∈ {0, 1}^*m*^, where:

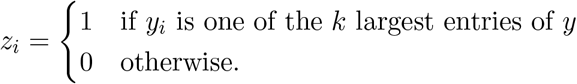

In other words, only the *k* neurons that fire at the highest rate among the population remain firing, and the rest are silenced. Mechanistically, this is implemented by inhibitory neurons, which receive excitatory input from the encoding neurons and provide feedback inhibition, which silences all except the highest firing neurons. This computation is often dubbed a “*k*-winners-take-all” competition [50–52].

Importantly, unlike the random hash functions typically used in count sketches, where a small change in the input could result in an arbitrarily far apart representation, this neural hash function is *locality-sensitive* [53–56]. This means that the more similar two inputs are, the more overlap there will be in their *z* representations. Biologically, this property is useful because it allows count estimates to be noise-tolerant [1]. In other words, instead of counting the frequency of 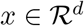, we want to count the total frequency of all items within a small radius around x, where the radius encapsulates noisy observations of *x*.

#### Synapse weight updating

The second piece involves modifying the synaptic weights *w* = (*w*_1_, *w*_2_, …, *w_m_*) of the *m* encoding neurons each time an input is observed. To mimic the way counters are updated in the count sketch, all weights are initialized to 0, and the update rule is:

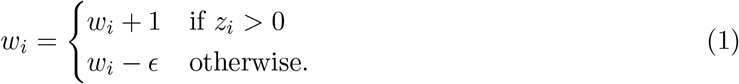

In other words, *w_i_* increases by 1 if *z_i_* is active for the input, and otherwise, *w_i_* remains the same, modulo a small memory decay parameter *ϵ* (in our experiments, we set *ϵ* = 0). This is effectively a Hebbian model (i.e., repetition enhancement) and leads to neurons whose activity scales with stimulus familiarity.

#### Frequency decoding

The third piece involves a read-out neuron, which outputs stimulus-specific frequencies. For a given input *x*, this neuron computes:

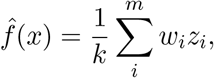

that is, the average of the *k* synapses activated for *x*, which is an estimate of the count of *x*. Since it may not be possible for a neuron to compute the average of its inputs, a simple alternative is to change the weight update in Equation (1) to *w_i_* = *w_i_* + 1/*k*, and then the decoder only needs to take the weighted sum of its inputs.

Thus, a fundamental counting data structure has a simple neural correlate.

### Deriving a “1-2-3-many” count sketch

While the neural circuit described above implements a count sketch data structure, there are several problems with this model in terms of neural plausibility. First, in computer science, count sketches are primarily designed to identify “heavy hitters” — i.e., very popular items, such as videos that are watched many times — with less precision in the counts of rare items. However, biologically, “light hitters”, such as items never seen before or just seen once or twice, are critical to distinguish because they signify novelty and degrees of familiarity. Second, behaviorally, the granularity of counts is likely not very high; e.g., it may not be possible (or even valuable) for organisms to distinguish between items seen 47 vs. 48 times, or between items seen 47 vs. 59 times. This is due to limits in the number of discrete firing rates that can be interpreted downstream as distinct, and limits in synaptic precision [57]. Third, experimental evidence suggests that recognition memory is largely based on repetition suppression [8–10, 58–62], as opposed to repetition enhancement.

To address these issues, we propose a “1-2-3-many” sketch, that only distinguishes between four categories of counts:

- **‘1’:** novel (first experience).
- **‘2’:** weakly familiar (more than just one random experience).
- **‘3’:** moderately familiar
- **‘many’:** strongly familiar (constantly re-occurring experiences)

We hypothesize that these four categories provide the best “bang for the buck”, in terms of ethological value to survival and precision to encode, with larger counts having increasingly diminishing returns. Novel items (category 1) are clearly important, as they alert organisms to new and potentially salient events [63]. However, many stimuli are experienced once (e.g., randomly), without much significance, and only a fraction of these stimuli are experienced twice. The two latter categories further help to separate environmental patterns from expected environmental stochasticity (Discussion). Thus, associating stimuli with graded levels of familiarity [58, 64, 65] could increase the behavioral repertoire of organisms.

How can we devise a 1-2-3-many sketch? The only change required is in the weight update rule. Previously, we initialized weights to 0 and applied a Hebbian update. Here, we initialize weights to 1 and apply an anti-Hebbian update, with the following functional form:

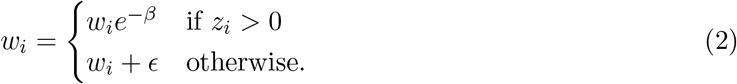

In other words, the weight is roughly 1 if the item is being experienced for the first time; *e*^−*β*^ for the second experience; *e*^−2*β*^ for the third experience; and less than *e*^−3*β*^ for all subsequent experiences.

Thus, novel items have large responses, which decrease multiplicatively with familiarity [59, 66], and the decoder neuron only needs to have four distinct responses, each representing a count category. Compared to the additive Hebbian model, this model creates greater separation between count categories, which makes it easier to read-out and control behavior (Discussion), at the expense of encoding fewer categories. In addition, all weights will be bounded between 0 and 1 (assuming *ϵ* = 0; otherwise, saturation can clip weights at 1).

### The neural count sketches accurately track item frequencies in streaming data

We tested the accuracy of count estimates from the two neural count sketches using streaming data from synthetic and real-world datasets, to demonstrate how well they work in practice.

#### Datasets and experimental setup

The first dataset, Synthetic, consists of *N* = 1000 items with *d* = 50 dimensions per item, where each dimension is drawn randomly from an exponential distribution. This distribution was selected because several types of neural stimuli, such as faces [67] and odors [51], are encoded as an exponential distribution of firing rates over a population of neurons. The second dataset, Odors, is experimentally collected response data of *d* = 24 olfactory receptor neurons in the fruit fly to *N* = 110 odors [68]. The third dataset, MNIST, consists of *N* = 10000 images of handwritten digits, where each image is of dimension *d* = 84. We reduced each dataset such that there were no pairs of items that were highly correlated (Pearson *r* ≥ 0.80) with each other. We did this because correlated items have highly overlapping representations and thus counts that would interfere with each other; moreover, such pairs of stimuli may be difficult for animals to distinguish without training. Nonetheless, many pairs of moderately correlated items were retained. For all datasets, we set *m* = 10000 (number of encoding neurons) and *k* = 10 (sparsity of the representation).

To generate the sequence of observed items, from each reduced dataset 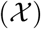, we drew *n* random samples with replacement according to a Zipf (power-law) distribution. The Zipf distribution captures frequency occurrence data in many domains [69], and allows us to explore the full gamut of counts, from those items never observed in the sequence to those observed many times.

After the *n* items were inserted into the sketch, we iterated through each unique item *x* in the dataset and compared its ground-truth count to its predicted count, 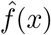, from the sketch. To test robustness to noise, we compared the ground-truth counts for *x* to the predicted count 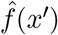, where *x*′ is the same as *x* but where each dimension is multiplied independently by a random value in [0.85,1.15] (i.e., up to 15% noise is added to *x*).

See Supplementary Methods for full details.

#### The Hebbian neural count sketch generates signals that scale with item frequencies

Recall that the neural count sketch uses a Hebbian learning model (i.e., repetition enhancement), and the output from the decoder neuron should correlate with the frequency of the item. This mimics neurons that become more active with familiarity.

On the Synthetic dataset, the output from the decoder neuron was highly correlated with the true count estimate (*r* = 0.935; Figure 2A). Without noise, all count estimates are either on or above the *y* = *x* line, since the count sketch is a biased estimator (i.e., it can over-estimate counts, but not under-estimate). With noise added (Figure 2B), the correlation only reduced to *r* = 0.880. Thus, count estimates for an item are robust to reasonable levels of variation in the item. This is due to the use of a locality-sensitive hash functions, which ensure that very similar items are mapped to overlapping representations in high dimensions [53–55].

**Figure 2:**
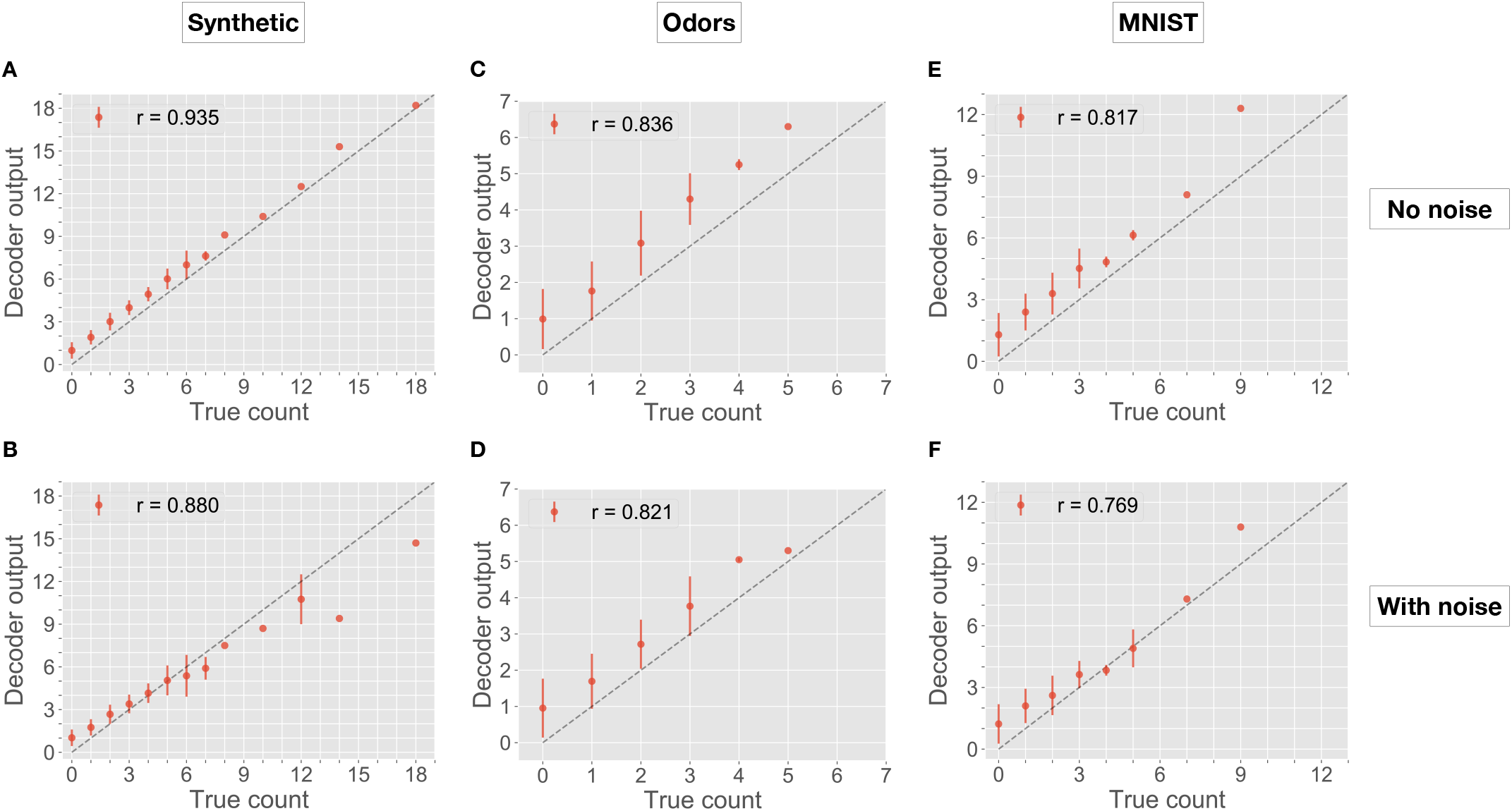
Performance of the neural count sketch (Hebbian model). In each panel, the *x*-axis shows the ground-truth count, and the *y*-axis shows the predicted count, as outputted from the decoder neuron. Dots show average decoder responses for items of the same ground-truth count, and errors bars indicate standard deviation. Perfect performance would lie on the dotted *y* = *x* line. Pearson correlation coefficient (*r*) quantifies performance accuracy (larger is better). Each panel shows a dataset (columns), with or without noise added to items (rows).

On the Odors and MNIST datasets, we observed similar trends, with a high correlation (*r* = 0.836 and *r* = 0.817; Figure 2C,E) between ground-truth and predicted counts without noise, and with small loss in performance with noise (*r* = 0.821 and *r* = 0.769; Figure 2D,F). Much of the error can be attributed to groups of moderately correlated items, whose counts collectively interfere with each other. For example, if we reduced the Odors dataset further by ensuring that the maximum pairwise similarity between any two items was *r* = 0.70 (instead of 0.80), then with noise, the correlation between predicted and true counts increases from 0.821 to 0.880.

Overall, the neural (Hebbian) implementation of the count sketch data structure works well in estimating counts, even for items that partially overlap.

#### The anti-Hebbian neural count sketch provides a mechanism to distinguish 1-2-3-many

Recall that the 1-2-3-many sketch uses an anti-Hebbian learning model (i.e., repetition suppression). To gauge performance of this sketch, we asked how distinguishable are the responses from the decoder neuron for items in the four count categories.

On all three datasets (Figure 3, top), we see characteristic repetition suppression, where novel items have large decoder responses, which are reduced with familiarity. For example, for the Odors dataset, items in category ‘1’ (novel) have an average response of 0.749 ± 0.168, whereas items in category ‘2’ have an average response of 0.351 ± 0.077, and this continues further with familiarity: 0.142 ± 0.034 for category ‘3’, and 0.040 ± 0.022 for ‘many’. All three comparisons — 1-vs-2, 2-vs-3, and 3-vs-many — are significantly different (*p* < 0.01; Wilcoxon rank sum test). With noise (Figure 3, bottom), there is more variation as expected, but all four categories remain distinguishable.

**Figure 3:**
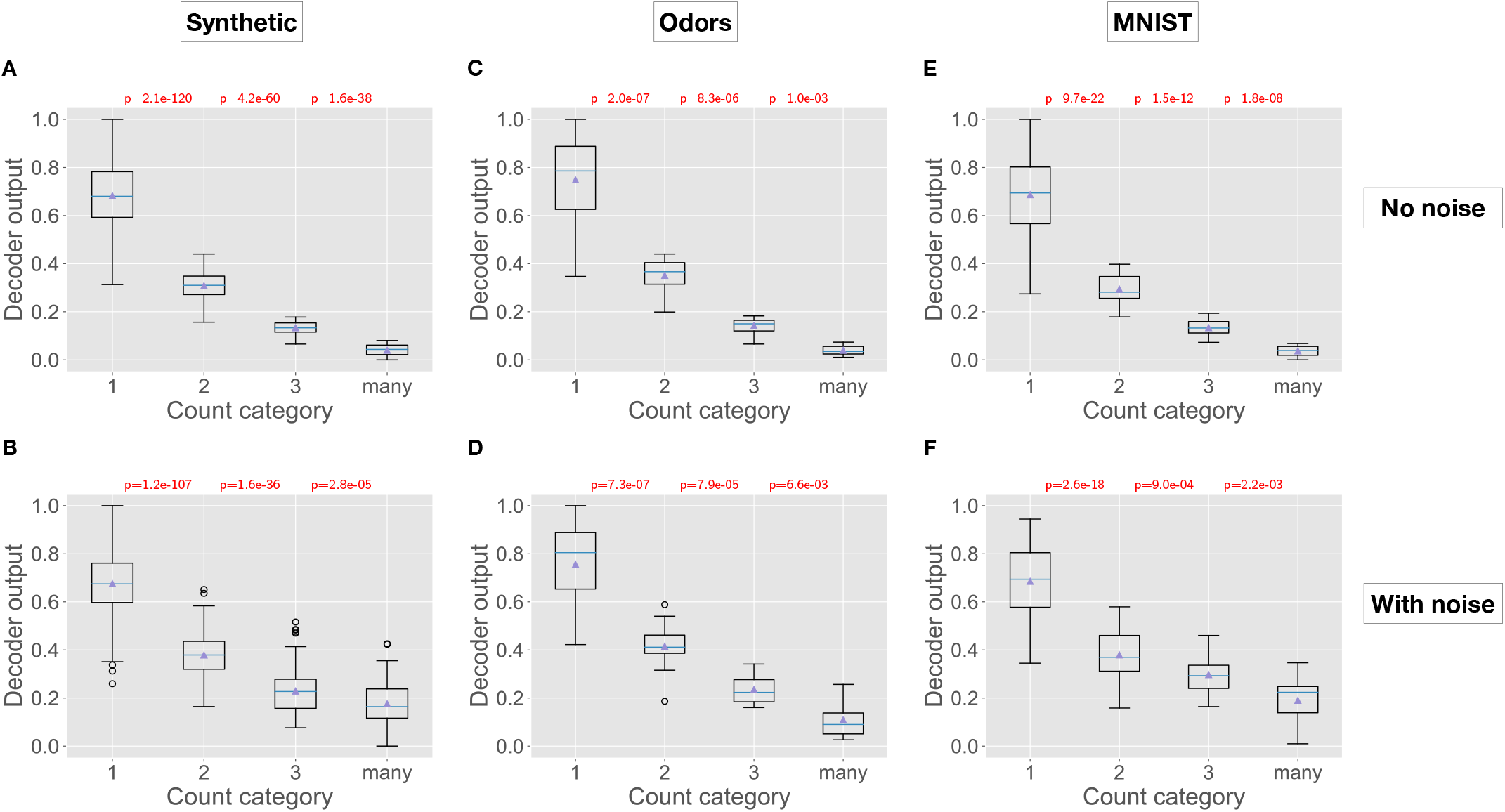
Performance of the 1-2-3-many sketch (anti-Hebbian model). In each panel, the x-axis shows the four ground-truth count categories, and the y-axis shows the decoder output from the 1-2-3-many sketch. For example, count category ‘1’ includes all items being observed for the first time (i.e., items not present in the input sequence), and the decoder output shows the response magnitude of the decoder neuron for all such items. Boxplots show the median and first and third quartiles, with whiskers extending from the box by 1.5-times the inter-quartile range. Shown at the top are *p*-values (Wilcoxon rank sum test) comparing differences in decoder response magnitudes between items in successive count categories. For example, in panel A, the difference between decoder responses to items in count category ‘1’ vs ‘2’ was statistically significant, with *p* = 2.1e-120. All *p* < 0.01 are colored red. Each panel shows a dataset (columns), with or without noise added to items (rows).

Thus, across three diverse datasets, the 1-2-3-many sketch provides sufficient granularity to robustly categorize items into four count categories.

### Theoretical analysis of the neural count sketches

To extrapolate from the empirical results and quantify how the accuracy of count estimates depend on environmental and neural circuit variables — such as the number of stimuli observed, the number of encoding neurons, the sparsity of representations, and synaptic precision — we mathematically analyzed the neural count sketch (Hebbian model) and the 1-2-3-many sketch (anti-Hebbian model). We summarize our main results below; full details and proofs can be found in Supplementary Information.

The primary setting we consider is one in which there are *N* distinct items (e.g., odors) that are *well-separated* from each other, in the sense that the distance between them is roughly what would be expected if they were chosen independently at random; this is formalized in Assumption 1. The sketching scheme is shown a sequence of *n* observations drawn from these *N* items, where the items are interleaved arbitrarily and might appear multiple times. Information about the observations gets coded in the weights *w_j_*, and when a subsequent query *x* (also one of the *N* items) is made, the sketch produces a frequency estimate for it. We study how close this frequency estimate is to the actual number of times *x* appeared in the sequence. All bounds hold with probability 1 – *δ*, where the confidence parameter 0 < *δ* < 1 impacts the manner in which *k* and *m* must be set.

For the neural count sketch, we prove (Theorem 2) that frequencies upto a value *f* are estimated within ±1 if the number of encoding neurons, *m* = *O*(*kn*), and if the sparsity, *k* = *O*(max(*n, f*^2^)log(1/*δ*)). For the 1-2-3-many sketch, we prove (Theorem 5) that it is sufficient to have *m* = *O*(*kN*) and *k* = *O*(log(1/*δ*)), which improves upon the neural count sketch in two important ways. First, the bound depends on the number of distinct items (*N*), rather than the total number of observations including repetitions (*n*), which could be far larger. Second, a significantly smaller setting of *k* (and thus *m*) is sufficient. In other words, the 1-2-3-many sketch only needs a few synapses to be allocated per unique item to generate good estimates.

The superior performance of the 1-2-3-many sketch comes at the cost of a higher weight precision requirement. The count sketch can accurately report frequencies upto *f* as long as its synaptic weights *w_j_* have *O*(log *f*) bits of precision. The 1-2-3-many sketch, on the other hand, needs *O*(*f*) bits of precision per weight, which is still within empirical estimates for small *f* (e.g., 3–5 [57]).

We also look at what happens when items are not necessarily well-separated. In such situations, where items lie in a continuum without well-defined boundaries, the notion of frequency becomes murkier. We show that nonetheless, the count sketch functions as a *kernel density estimate* [70], where the sketch outputs a value that relates to the density of observations around a given item. Kernel density estimators serve as a foundation for numerous machine learning problems that require understanding the “shape” of the underlying data (Discussion).

Overall, these mathematical proofs provide bounds on how accurately stimuli can be tracked using the two neural count sketches.

### The *Drosophila* mushroom body implements the anti-Hebbian count sketch

Here, we provide some evidence supporting the “1-2-3-many” model from the olfactory system of the fruit fly, where circuit anatomy and physiology have been well-mapped at synaptic resolution [71, 72]. The evidence described below includes the neural architecture of stimulus encoding, the plasticity induced at the encoding-decoding synapse, and the response precision of the decoding (counting) neuron. The latter two we derive from a re-analysis of data detailing novelty detection mechanisms in the fruit fly mushroom body [8], where odor memories are stored.

#### Stimulus encoding (Figure 4A)

**Figure 4:**
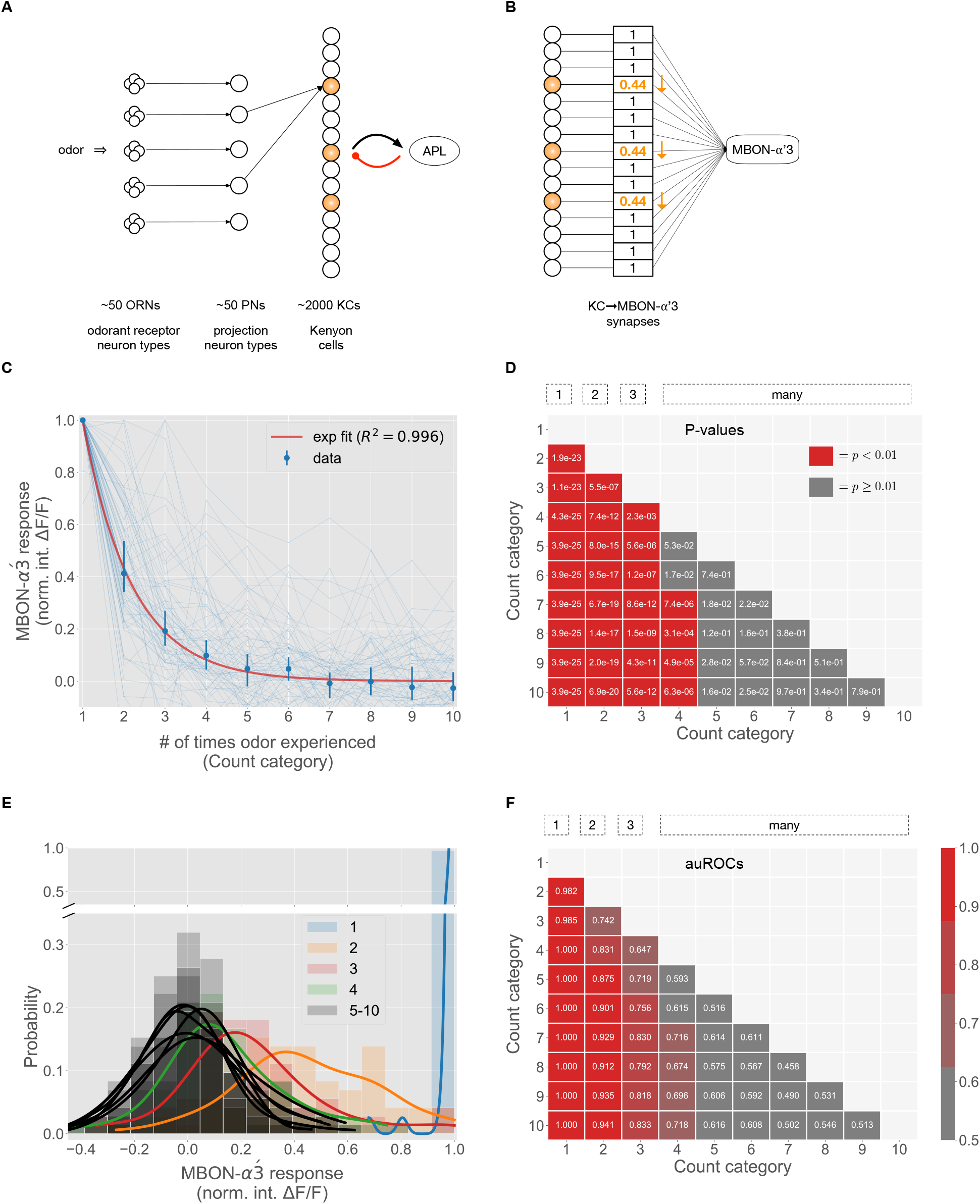
Experimental evidence of the “1-2-3-many” sketch from the insect mushroom body. **A)** Schematic of the fruit fly olfactory system. Odors are initially represented by the firing rates of 50 odorant receptor neuron types, which send axons to 50 projection neuron (PN) types. PNs then send odor information to 2000 Kenyon cells (KCs), each of which provides feed-forward excitation to a large inhibitory neuron (called APL), which sparsifies the KC representation via feedback inhibition. **B)** Synapses between activated KCs and the counting neuron (MBON-*α*’3) are modified (weakened) when an odor is experienced. **C)** Response dynamics of MBON-*α*’3 (y-axis) over 10 successive presentations of odor MCH (*x*-axis). Data shows responses of 72 cells (light blue) over 59 flies. Blue dots (dark blue) show median response values, and error bars show 99% confidence intervals determined by 20,000 bootstraps. For each cell, responses are normalized to the magnitude of the first presentation. Red curve shows data fit to an exponential function (*y* = *ae^bx^*), with a suppression constant of 0.44. **D)** Heatmap of *p*-values (Wilcoxon rank sum test) comparing differences in response magnitudes for all pairs of count categories. For example, MBON-*α*’3 responses are significantly different comparing an odor seen for the first time vs. the second time (*p* = 1.9e-23), but responses are not significantly different comparing the 4th vs. the 5th experience (*p* = 5.3e-02). Red blocks indicate *p* < 0.01, and gray blocks indicate *p* ≥ 0.01. **E)** Histogram distributions of MBON-*α*’3 responses for each count category, with kernel density estimation curves plotted on top. Categories 1, 2, 3, and 4 are shown in blue, orange, red, and green, respectively; categories 5–10 are shown in black. Categories 4–10 (many) are highly overlapping and distinguishable from categories 1, 2, and 3. **F)** Heatmap of area under the ROC (auROC) values for discriminating between each pair of count categories using a linear threshold. For example, the auROC for discriminating 1-vs-2 was 0.982. Distinguishability is highest for categories 1, 2, and 3 (first three columns), and then tapers off for subsequent categories.

In the fruit fly olfactory system [73], odors are initially represented by the firing rates of *d* = 50 types of odorant receptor neurons. After a series of pre-processing steps, including gain control [74, 75], noise reduction [76], and divisive normalization [51, 77], odors are represented by the firing rates of *d* = 50 types of projection neurons (PNs), which each receive input from sensory neurons expressing the same receptor type. Thus, an odor *x* is a point in 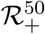.

The first piece (assigning the odor a sparse, high-dimensional representation) is accomplished by 2000 Kenyon cells (KCs), which receive input from the PNs. Each KC samples randomly from approximately 6 of the 50 PN types [78] and sums up their firing rates. Hence, the random projection matrix *M* is a sparse binary matrix, with about 6 ones per row. Next, each KC sends feed-forward excitation to an inhibitory neuron, called APL, which then sends feed-back inhibition to each KC. As a result, only the top 5% of highest-firing KCs remain active for the odor, and the rest are silenced [45, 50, 51]. Moreover, KCs tend to respond in a binary manner, firing either zero spikes or just a few spikes per odor [45, 79, 80]. Thus, odor are encoded as a high-dimensional vector (with dimension *m* = 2000), of which only a few KCs (*k* = 100) are active for the odor.

#### Synapse weight updating (Figure 4B–C)

The second piece involves synaptic connections from KCs to an output neuron. In the fly mushroom body, there are 35 types of output neurons (called MBONs [72, 81]) that read-out information from the 2000 KCs and control behaviors, such as learning to approach or avoid odors [73]. KC→MBON synapses are plastic [82], and dopamine modulates the synaptic strength bi-directionally depending on the timing contingency between KC activity and dopamine release [8, 83, 84] in a manner consistent with anti-Hebbian plasticity (albeit on a longer time scale than traditional STDP and without requiring post-synaptic firing [85]).

Recently, one such MBON (called MBON-*α*’3) was discovered that computes the novelty of an odor [8] (Figure 4B). When an odor is experienced, synapses from the odor’s activated KCs onto MBON-*α*’3 multiplicatively weaken, whereas synapses from non-active KCs onto MBON-*α*’3 strengthen slightly (*ϵ* in Equation (2)). The output of MBON-*α*’3 is the weighted sum of its inputs (i.e., the activity of each KC multiplied by its synaptic strength). Thus, repeated exposure to the same odor depresses active KC→MBON-*α*’3 synapses, which suppresses the activity of MBON-*α*’3 in response to the odor, indicating that the odor has become familiar. Hattori et al. [8] also found another output neuron (called MBON-*β*1>*α*) that responds linearly with familiarity. Thus, this circuit uses repetition suppression (MBON-*α*’3 for novelty) and possibly repetition enhancement (MBON-*β*1>*α* for familiarity), though the latter remains unconfirmed mechanistically.

To quantify the weakening in the KC→MBON-*α*’3 synaptic weights following stimulus experience, we re-analyzed MBON-*α*’3 responses from 72 cells to 10 repeated exposures of the same odor (Figure 4C). Each exposure increases the number of times the odor is experienced. The median response of MBON-*α*’3 to an odor experienced for the first time (category 1) was 1.00, compared to 0.413, 0.193, 0.098, and 0.048, for categories 2 through 5, respectively. The data closely fit an exponential decay function (*R*^2^ = 0.996), with a suppression constant of 0.44. This means that each successive exposure decays the MBON-*α*’3 response by a factor of 0.44. Thus, *β* = – ln(0.44) in Equation (2), supporting the general functional form of suppression proposed.

#### Frequency decoding (Figure 4D–F)

While MBON-*α*’3 was originally conceived as a binary novelty detector neuron [8], our re-analysis of MBON-*α*’3 responses provides evidence for the presence of more than two count categories along the novelty-familiarity axis. To show this, the activity level of MBON-*α*’3 must be significantly different across multiple experiences of the same odor. At some point, the difference in activity between successive experiences becomes indistinguishable, and this is where the “many” category kicks in, indicating that responses to all subsequent experiences are essentially the same. Specifically, for “count category” *j* to exist, it must be possible to distinguish category *j* from each other category, including each individual category encapsulated by “many”.

Strikingly, re-analysis of MBON-*α*’3 activity levels to successive experiences of an odor shows that the distinguishability of responses are consistent with the 1-2-3-many model (Figure 4D). Categories 1, 2, and 3 were each significantly different from each other category (all *p* < 0.01; Wilcoxon rank sum test). However, category 4 was not significantly different from categories 5 and 6, and categories *j* = 5 onwards were not significantly different from categories *j* + 1 onwards. Thus, the decoding neuron can robustly distinguish among odors experienced 1, 2, or 3-times before, with a separate category for 4 or more (many).

Visualization of the distributions of MBON-*α*’3 responses to odors in each count category shows the separability of categories 1, 2, and 3, as well as the clustering of categories 4–10 (Figure 4E). The blue curve (category 1) is clearly distinguishable from the orange curve (category 2), which is distinguishable from the red curve (category 3). However, the curves for categories 4 (green) and 5–10 (black) are highly overlapping, indicating that their responses are roughly the same and comprise the ‘many’ category. We also quantified the separability of all pairs of count categories using a simple response threshold discrimination model (Figure 4F). The area under the ROC curve remained high (≥ 0.70) when discriminating between 1, 2, and 3 and nearly all other categories, but was considerably degraded for subsequent categories, further supporting the existence of four robust count categories.

These results suggest that MBON-*α*’3 encodes frequency information about odor memories into four distinct categories along the novelty-familiarity axis, which may be used downstream in the circuit to modulate behavior.

## Discussion

### Summary

One role of theory in neuroscience is to propose plausible circuit mechanisms that support important neural computations. Here, we showed how a fundamental data structure used by computer scientists to count frequency events in streaming data could be implemented by canonical neural circuitry. This theory was supported by experimental data in the insect mushroom body, which gave credence to the 1-2-3-many count sketch, both qualitatively and quantitatively, in terms of the required neural architecture, the functional form of synaptic plasticity, and the output precision of the counting neuron.

Our proposed neural count sketch data structure has four properties: (i) it provides counts that are stimulus-specific; (ii) it has a large storage capacity, that is, it requires only a few synapses per unique item [18]; (iii) it offers robustness, that is, the ability to generalize counts across noisy versions of the same item; and (iv) it is fast and automatic, providing frequency estimates of inputs after two synapses of computation, requiring only tens to hundreds of milliseconds.

### Experimental questions and testable predictions

Our work raises several experimental and circuit design questions.

First, how might downstream mechanisms robustly read-out frequency estimates and use them to modify behavior? For the anti-Hebbian model, this would require interpreting the four firing rates of the 1-2-3-many counting neuron as distinct. One option is to convert this continuous firing rate into a discrete (i.e., a “one-hot” encoded) representation (Figure 5A). For example, the counting neuron could synapse with four output neurons, each with successively lower firing thresholds and with inhibition from neurons with higher thresholds to neurons with lower thresholds. As a result, each count category will be represented by the activity of a single neuron. A second option is to hierarchically string together counting neurons (Figure 5B). Here, one counting neuron inhibits the activity and synaptic plasticity of another counting neuron, such that the first neuron robustly encodes 1 and 2, and (after the inhibition from the first neuron is lifted), the second neuron encodes 3 and many, etc. This option provides a mechanism to translate a small resolution counting system to a larger one, with greater separability between count categories. Thus, multiplexing counting modules via hierarchical connections could provide robustness and scalability.

**Figure 5:**
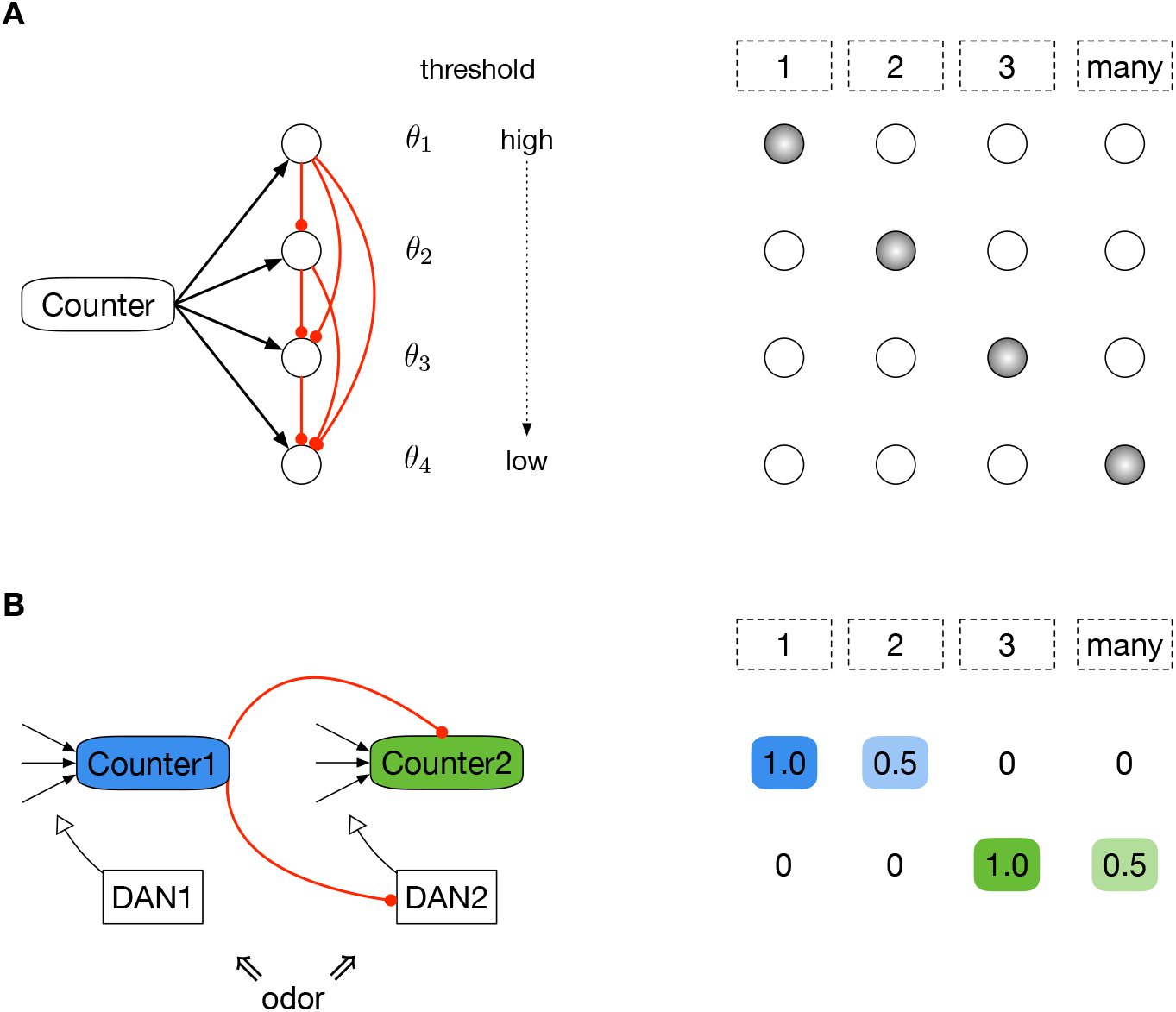
Hypothetical read-out mechanisms of the counting neuron. **A)** This model translates the continuous firing rate of the 1-2-3-many counter neuron into a discrete representation. The counter synapses onto four downstream neurons, each with succcessively smaller firing thresholds (*θ*_1_ > *θ*_2_ > *θ*_3_ > *θ*_4_). Neurons with larger thresholds inhibit those with smaller thresholds. As a result, each count category becomes “one-hot” encoded, making it easier to modify behavior. **B)** This model hierarchically strings together counting neurons to increase scalability and the resolution between count categories. In the insect mushroom body, odors activate dopamine neurons (DANs), which modulate synaptic weights onto counting neurons. In this model, there are two counting neurons with two associated DANs. Both counters receive input from encoding neurons, and Counter1 inhibits Counter2 and DAN2. Counter1 encodes categories 1 and 2 with high (1.0) and medium (0.5) responses, respectively. When Counter1’s activity is diminished after the second stimulus experience, the inhibition is lifted, allowing Counter2 to encode the two subsequent categories with high and medium responses.

For the Hebbian model, the read-out may simply be the total activity level, which scales with stimulus frequency in a continuous manner. Indeed, in the mushroom body, the response of the familiarity neuron (MBON-*β*1>*α* [8]) increases linearly with successive odor experience, which supports the additive form of synaptic plasticity in Equation (1). Alternatively, a discrete read-out could be generated by applying a sigmoid activation function to the counting neuron. Category 1 would correspond to the response prior to the rise of the sigmoid, with a few categories in the middle, and then ‘many’ at the saturation of the sigmoid.

Second, our results suggest that behaviorally, animals can distinguish among stimuli in each of the four count categories, as opposed to just the traditional novel vs. familiar categorization. Ethologically, it seems important for organisms to discriminate between the first and second experience of a stimulus, since there are many things experienced once (e.g., randomly) but many fewer things experienced twice. Distinguishing between the second and third experiences may be advantageous during exploratory behavior. For example, an animal might enter and then leave a locale with some identifying scent, experiencing it twice, once upon entry and once more upon exit; returning again to the same locale could trigger a memory that the animal has already been there before. Similarly, another animal (say, a potential mate) may enter and then leave a locale, and knowing if that animal returns again could warrant a change in behavior. Indeed, many things come and go, but few things come back again. The final category hosts stimuli experienced ‘many’ times, indicative of re-occurring experiences that define one’s environment (e.g., a mother’s voice, the scent of a nest). It is also striking that some indigenous tribes only have words for “one”, “two”, “three”, and “many” [86], which suggests that the value of having four distinct count categories may indeed be broadly conserved, even in humans.

Third, we analyzed the functional form of repetition suppression at single cell resolution, and we quantified how the setting of *β* (the suppression constant) and other circuit parameters impact the distinguishability of count categories (Theorem 5). How general is this form and the corresponding value of *β* in the numerous other systems that use repetition suppression to encode stimulus familiarity [9, 10, 58–62]? Our theory also hypothesizes that count estimates are privy to the similarity structure of stimuli. For discrete, well-separated stimuli, our model predicts that animals can generalize counts across noisy versions of the same stimuli. For continuous stimuli, count estimates may reflect a kernel density estimate, capable of counting sub-features shared by stimuli.

Fourth, what are the factors, such as attention [87], arousal, and other brain states [83, 88, 89], that control whether counts are updated upon stimulus experience? In the mushroom body, repetition suppression occurs due to dopamine release in the *α*’3 compartment after each experience of a stimulus. The lack of dopamine release may be indicative of an experience that is not “inserted” into the sketch and hence not remembered. This mechanism also provides the intriguing benefit of being able to query the count sketch for the frequency estimate of an item, without updating its count — i.e., a form of “recollection”. In addition, the unit of “experience” that triggers dopamine release remains unclear. For images, is a single 2-second exposure equivalent to five successive exposures of 400ms each? For odors, what duration of an odor puff gets integrated into a single experience?

Fifth, what is the function of the many other “counting neurons” in the brain that track stimulus familiarity? One idea is that counts are conditioned on location; e.g., “how many times have we met in New York?” The hippocampus is believed to be a central location where counts and context may be integrated [2, 9, 90–92]. Another idea is that some neurons have faster or slower synaptic recovery rates (*ϵ*), and thus, different memory spans. For example, in the insect mushroom body, different anatomical compartments acquire and forget memories at different rates, leading to short- and long-term memories [93]. For counting, non-zero values of *ϵ* provide a mechanism to free-up capacity for newer items at the expense of those not experienced in a while. This would also help prevent synapse saturation (to 1 for the Hebbian model, and to 0 for the anti-Hebbian model). Relatedly, there are variants of count sketches that allow for item deletion [94, 95]. Thus, having multiple counting neurons can help contextualize frequency estimates across both space and time.

### Generality to other brain regions and species

There are two main ingredients of our data structure — sparse, high-dimensional representations for stimuli and repetition-based modulation of synaptic weights. Where else are these two features found in the brain? Sparse, high-dimensional representations are ubiquitous in sensory areas, such as in olfaction, vision, audition, and somatosensation, and in the hippocampus [42, 96]. Some of these regions also employ computations, such as decorrelation [97], sharpening [3, 64], and pattern completion, which would further boost the stimulus-specificity of counts. Repetition suppression has been observed in many mammalian brain regions, including the perirhinal cortex, prefrontal cortex, basal ganglia, and inferior temporal cortex, amongst others [9, 10, 98]. Repetition enhancement (e.g., familiarity neurons) have also been found in many of these regions [12, 99], though less common. Thus, all the machinery required to implement count sketches are prevalent in the brain.

### Applications to machine learning

In computer science, count sketches are typically used to identify “heavy hitters” (i.e., very popular content), which typically constitute a small fraction of the observed items. However, an equally important class of items are “light hitters”, that is, those items that are rare or have never been seen before, which may signal anomalies and require attention. The 1-2-3-many sketch serves as a bridge between these two extremes, by providing fine resolution at the transition between novel and familiar, as well as a separate class (“many”) for popular items. For example, in reinforcement learning, rewards are often sparse, and the novelty-familiarity spectrum can be supplemented as an intrinsic reward signal to drive exploration and the discovery of desirable states [1]. In addition, number neurons were shown to spontaneously emerge in artificial neural networks trained on object recognition tasks [100], suggesting that pre-loading deep networks with computational modules for frequency estimation may be a useful component towards generalized decision-making.

## Acknowledgements

The authors thank Alison L. Barth, Tatiana Engel, David Freedman, Partha Mitra, Guruprasad Raghavan, Yang Shen, and Shyam Srinivasan for helpful discussions.

## Notes

### Competing Interest Statement

The authors have declared no competing interest.

